# APP Dosage and Extracellular Domain Variants Drive Distinct Defects in Neurogenesis modeled in Down Syndrome iPS cells

**DOI:** 10.64898/2026.08.05.742240

**Authors:** Melvys Valledor, Kelly Smith, Jeanne B. Lawrence, Lucas J. Sosa

## Abstract

The amyloid precursor protein (APP) is heavily studied as the source of amyloid beta in Alzheimer’s disease (AD), however, the complex functions of APP remain poorly understood, as does the impact of APP dosage on neurodevelopment. Here we study APP specifically in the context of Trisomy 21. In an effort to reduce APP dosage in trisomy 21 iPSCs, we generated trisomic isogenic lines which vary in APP dosage, including a full APP knock-out line, as well as lines carrying mutations of the APP extracellular domain. We used a panel of these lines to study potential impacts of APP dosage or structure on two distinct steps of neurogenesis in trisomic cells: 1) terminal differentiation of human neuro-progenitor cells (NPC) to post-mitotic neurons and 2) neuron structure as reflected in neurite outgrowth. Complete loss of APP causes marked defects in each of these two distinct steps, reducing both the terminal differentiation of NPCs to neurons, and proper neurite development for extended neuron structure. Hence, APP is necessary for both aspects of normal neurogenesis. Further analyses of the null and other mutant lines indicate that APP likely impacts these two distinct steps by two different mechanisms. Collective results suggest that the reduced terminal differentiation of NPCs reflects an effect of APP dosage, whereas the defects in neurite extension are due to structural mutation of the APP extracellular domain. Absence of APP or reduced (monosomic) APP dosage prolonged the cycling of trisomic NPCs, which is known to be regulated by Notch signaling. APP and Notch are the main targets of gamma-secretase cleavage, hence we hypothesized that APP dosage may impact neurogenesis indirectly, potentially via effects on Notch signaling. To test this, we treated NPCs with Compound E which inhibits gamma-secretase (and Notch signaling); results show this restored levels of neurogenesis in APP depleted lines, supporting an indirect effect of APP *dosage*. In contrast, results indicate that disruption of APP extracellular domain integrity impacts neurite extension via a more direct role of APP in neuron structural maturation. This study describes a resource of well-characterized APP mutant isogenic DS iPSC lines, implicates a dynamic interplay between APP dosage and Notch signaling, and raises new questions about the impact of APP dosage in orchestrating neural progenitor fate decisions during human brain development, specifically in the context of trisomy 21.

## INTRODUCTION

The amyloid precursor protein (APP), encoded on Chromosome 21(Chr21), is generally important in pathogenesis of Alzheimer’s Disease (AD); APP triplication drives early onset AD in both Down syndrome (DS) and in APP microduplication syndrome (Fortea et al., 2020; Rovelet-lecrux et al., 2006). While APP is heavily studied in the context of AD, the broader biological roles of this widely expressed protein are still poorly understood (Coronel et al., 2018; Dunot et al., 2023; Müller et al., 2017; Sosa et al., 2017). Mouse studies have indicated that APP may have a role in the maturation and migration of cortical neurons (Callahan et al., 2017; Peralta Cuasolo et al., 2023; Young-Pearse et al., 2007) as well as neurite-axonal outgrowth, contact guidance, dendrite complexity (Sosa et al., 2017; Wang et al., 2016). Studies of APP potential role in neural progenitor cell (NPC) proliferation and differentiation using mouse models have yielded varying results (Bolós et al., 2014; Hu et al., 2013) reviewed by (Coronel et al., 2018; Dunot et al., 2023), although a recent study in euploid human iPS cells indicates that APP is required to balance the proliferation versus differentiation of neural stem cells (Shabani et al., 2023).

Many studies have examined APP biology in euploid systems, but here we investigate APP in the context of trisomy 21. Multiple lines of evidence in different sytems haveindicated that APP functions as a cell adhesion molecule which may support a role in neurite outgrowth and contact guidance, (reviewed in (Sosa et al., 2017)). However, i*n vitro* analysis of cultured mouse neurons from Ts65Dn (trisomic) mice showed evidence of perturbed neuronal neurite outgrowth and contact guidance and raised the possibility of an APP dosage effect (Sosa et al., 2014). However, interpretations are complicated by the additional complexity of trisomy 21 and potential differences between samples and/or hybrid mice background. Here, we compare neural cells differentiated from isogenic human trisomic iPSC lines with the same genomic background except for dosage and/or structural differences in APP. In addition to effects on neuron structure, we examine an earlier developmental step, the transition of cycling NSCs to post-mitotic neurons. In other work we have used a similar panel of isogenic DS iPSCs to investigate the collective effects of trisomy for all chr21 genes, using XIST-based “trisomy 21 silencing” during early neural differentiation (Czermiński & Lawrence, 2020); reviewed by (Gupta et al., 2024). Results showed that induced silencing of one chromosome 21 in otherwise identical cell cultures caused an increase in the terminal differentiation of NSCs to neurons. Single-cell RNAseq showed the prolonged cycling of neural stem cells and reduced neuron formation, which was linked to elevated Notch signaling (which drives cell cycling). This deficit in terminal differentiation was overcome by inhibition of gamma secretase which normally cleaves Notch to drive signaling. Importantly, both Notch and APP are the main cleavage targets of gamma-secretase (Zhang et al., 2000) and may have indirect interplay, such as via competition for cleavage by gamma secretase (Berezovska et al., 2001).

Having studied the effects of reducing over-expression of essentially all chr21 genes, here we manipulate just the APP gene in subclones of the same isogenic trisomic iPSCs. While our initial intent was to reduce APP dosage in trisomy 21 iPS cells, in pursuing this goal we generated a unique set of isogenic APP-depleted/mutated DS-iPSC cell lines, several of which we characterize in detail. This set of isogenic lines allowed us to investigate the impact of APP variants during neurodevelopment, including in cells fully depleted of APP. Our results indicate that APP protein impacts two crucial steps in early human neurodevelopment, as modeled here *in vitro*. Results suggest that APP *dosage* impacts the differentiation of cycling neural progenitor cells (NPC) into post-mitotic neurons. Even in APP null cells the reduced differentiation to form neurons can be overcome by gamma-secretase inhibition, which is known to impact Notch signaling and also APP cleavage. Rather than a direct functional role of APP required for terminal differentiation of NPCs, results suggest that APP levels may impact rates of neurogenesis indirectly, possibly by balancing or influencing Notch signaling in trisomic cells. In contrast, at a later step APP may be play a more direct role in supporting the structural maturation of neurons. In cells with two normal APP and one allele which skip exon 10 of the E2 domain, the truncated APP protein causes severe disruption of neuron structure, preventing normal extension of neurites. Consistent with a dominant negative effect on a required APP function, a very similar impact is seen in the APP null cells. Overall, results here indicate that APP dosage can indirectly impact the balance of NPC cycling versus neurogenesis, whereas APP plays a more direct functional role in neuron structure.

## RESULTS

### Generation and characterization of isogenic Down syndrome iPSCs lines with different modifications of APP

In order to reduce APP copy number in trisomic DS-iPSC cell lines (APP+/+/+), we used CRISPR/Cas9 to introduce double-strand breaks in exon 9 or exon 10 and co-transfected a donor construct carrying a selection cassette **(Figure 1A).** Cas9 cutting at APP produced many subclones that had undergone homology-directed repair to integrate the donor cassette, which were enriched by puromycin selection. With the initial goal of identifying cells with APP reduced to disomy, we then used two-color FISH, as shown in **Fig. 1B**, to screen for clones that had inserted the transgene (green) into one of three APP loci (red). In addition to the parental line we isolated and fully characterized three subclones which had inserted the transgene into just one APP allele, and we began to examine these lines for neural differentiation, as further described below. However, subsequent sequence analysis of all APP alleles revealed small indels in other alleles, as is known can occur using CRISPR in which breaks can be repaired by error prone non-homologous end joining. When we sequenced the APP locus in multiple clones, we found a combination of donor insertions and indels, and in the case of the donor-inserted exon 10 allele an unexpected in-frame exon 10 skipping event. In fact, from careful screening of the many subclones generated from this effort we found none with only a straightforward disruption of one APP allele. Nonetheless, our analysis yielded a panel of APP-edited lines ranging from complete APP knockout to reduced-dosage and E2-domain structural mutants as described in **(Fig. 1C)** and below.

**Figure 1.**
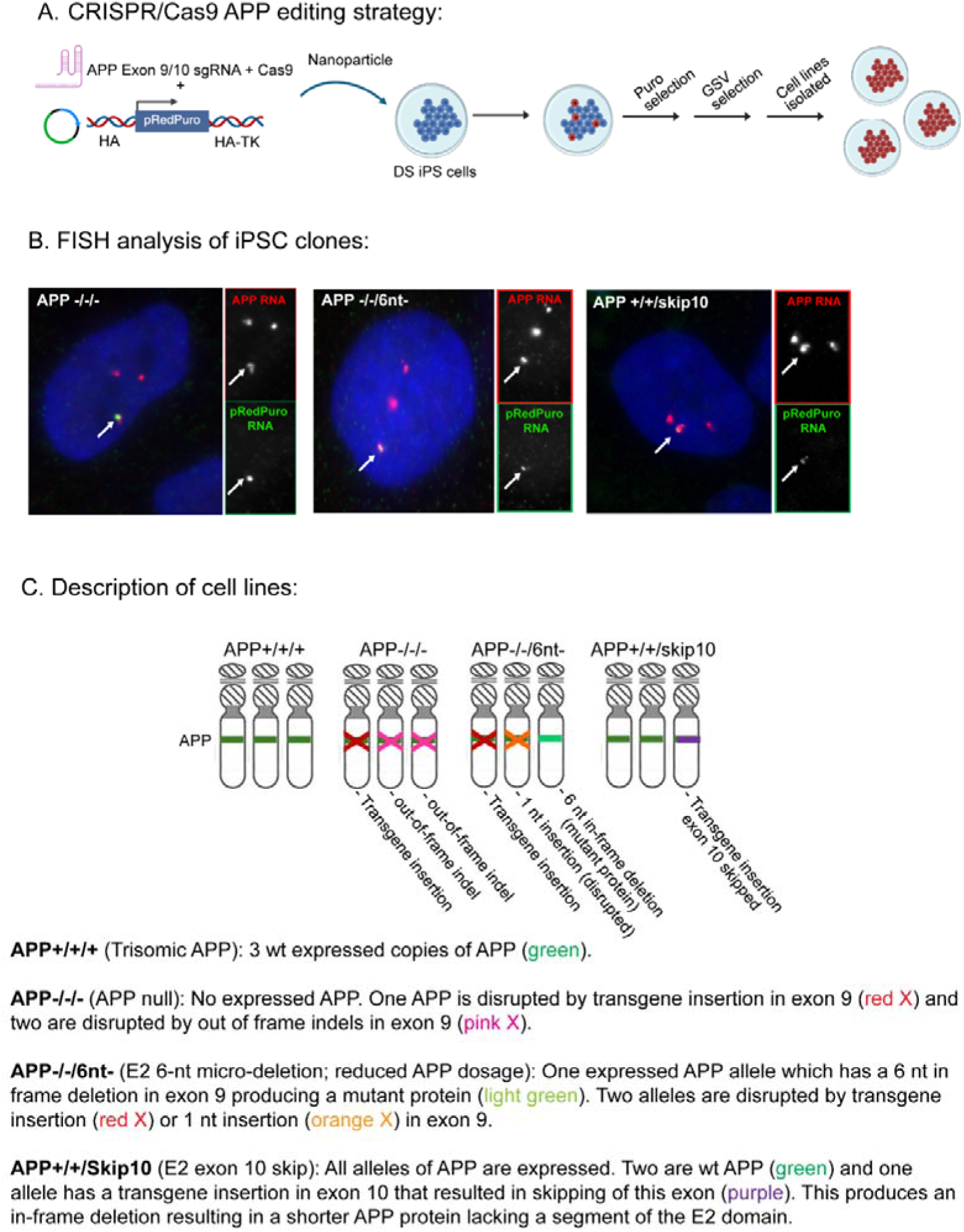
Generation of APP depleted DS-iPSC lines panel via CRISPR/Cas9-mediated gene disruption. **A.** Schematic representation of CRISPR/Cas9 genome editing strategy used to disrupt the APP locus in parental trisomic DS-iPSC cell lines (APP+/+/+). **B.** FISH analysis of 3 clones with targeted insertion of transgene (green) into an APP gene (red). **C.** Description of APP clones used in this study.

We summarize our analysis of the cell clones chosen for further study here. To show the correct integration site of the transgene, we performed PCR with primers that recognize the APP locus and the pRedPuro insert flanking regions **(Figure S1a)**. We observed expected products from the genomic DNA of sgRNA 1 (exon10) and sgRNA 4 (exon 9) corresponding to the DS-iPSCs cell lines we termed APP+/+/Skip10 and APP-/-/-, respectively **(Figure S1b-c).** To evaluate CRISPR efficiency and potential off target effects, we used a topo cloning and sequencing strategy. In the APP-/-/- cell line, CRISPR targeting induced indels at the other two APP alleles, causing a complete loss of APP in this cell line **(Fig. S2).**

**Figure 1C** summarizes the four APP genotypes we focused on for further analysis: (1) the parental trisomic DS-iPSC line, APP+/+/+, which carries three intact APP alleles and expresses APP at high levels; (2) an APP-null line, APP-/-/-, in which donor insertion and CRISPR-induced indels disrupt all three APP alleles, resulting in complete loss of APP expression; (3) an E2 domain mutant line, APP+/+/Skip10, which retains two intact APP alleles and has one donor-inserted in exon 10 allele; we discovered this does not lead to non-sense mediated decay put produces an in-frame exon-10-skip transcript encoding a shorter APP protein lacking the large exon-10-encoded segment of the E2 domain **(Figure S3)**; and (4) a reduced-dosage line, APP-/-/6nt-, in which APP expression is effectively monosomic and the single expressed allele carries a very small 6-nt in-frame deletion in exon 9 (p.Ala298_Val299del) at the N-terminal edge of the E2 domain **(Figure S3)**.

We characterized APP at the DNA, RNA, and protein levels in these isogenic DS-iPSC cell lines, with results summarized in **Table 1**. Western blot and immunofluorescence studies confirmed complete lack of APP protein expression in the APP-/-/- null line, and reduction in APP-/-/6nt- and APP+/+/Skip10 lines in comparison to the APP+/+/+ line **(Fig. 2)**.

**Figure 2:**
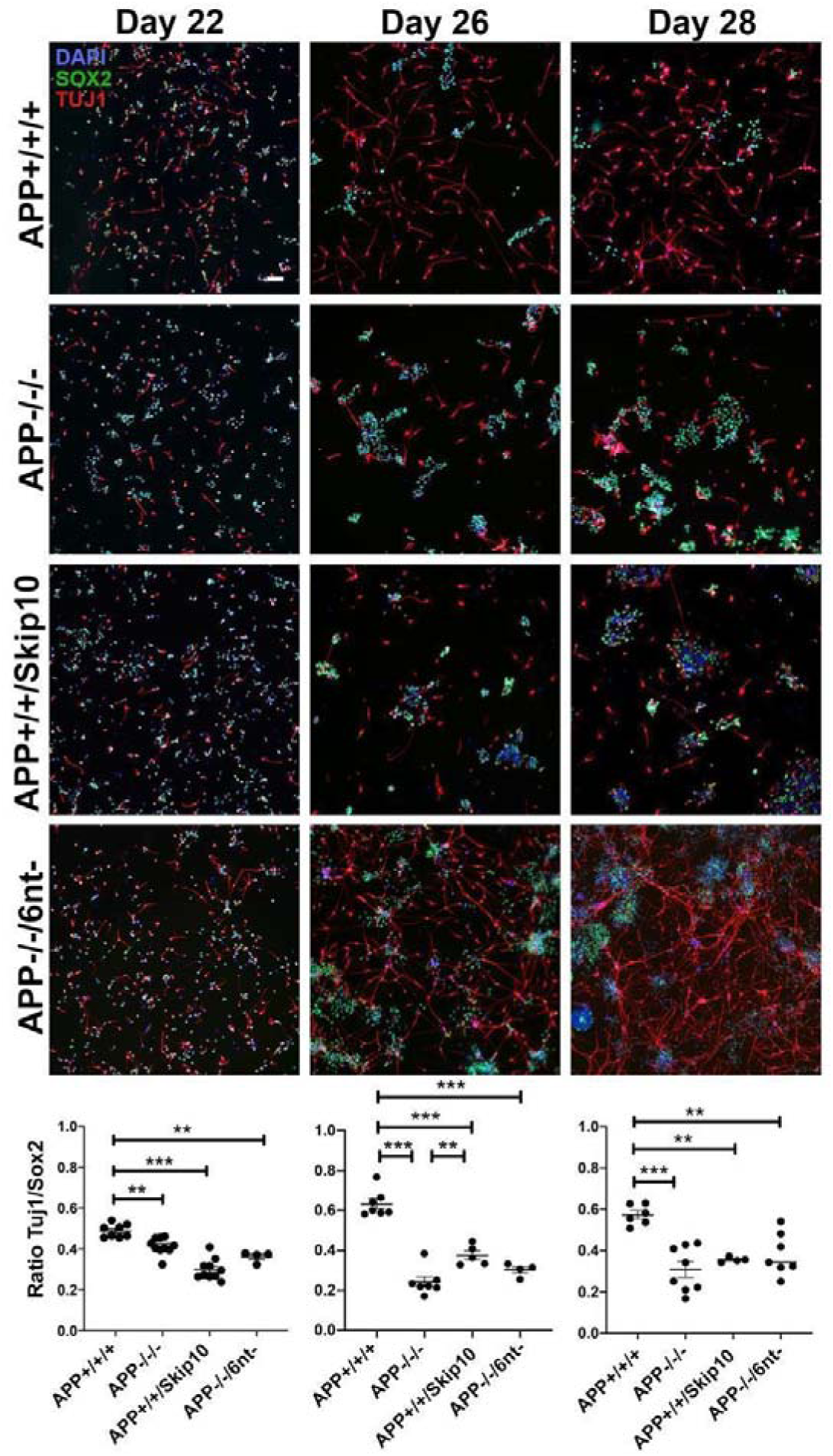
APP knock-out or structural mutation in isogenic lines reduces the terminal differentiation of NPCs into neurons. Representative immunofluorescence images stained for SOX2 (neural progenitor cells, green) and TUJ1 (neurons, red) of parental DS-iPSCs (APP+/+/+) and isogenic APP depleted lines (APP-/-/-, APP-/-/6nt- and APP+/+/Skip10) after neural differentiation for 22, 26 and 28 days. DAPI (blue) stain nuclei. APP-depleted/mutated lines exhibit reduced neuronal differentiation compared to APP+/+/+ controls. Scale bar, 50 μm. Graphs at bottom quantify the TUJ1⁺/SOX2⁺ cell ratio for the cell lines from 6 or more random low-magnification fields (represented by dots) in 1-2 samples for each of three time points. APP-depleted/mutated lines have a markedly reduced TUJ1⁺/SOX2⁺ ratio at later time points, reflecting impaired neuronal differentiation from cycling neural stem cells.

**Table 1.**
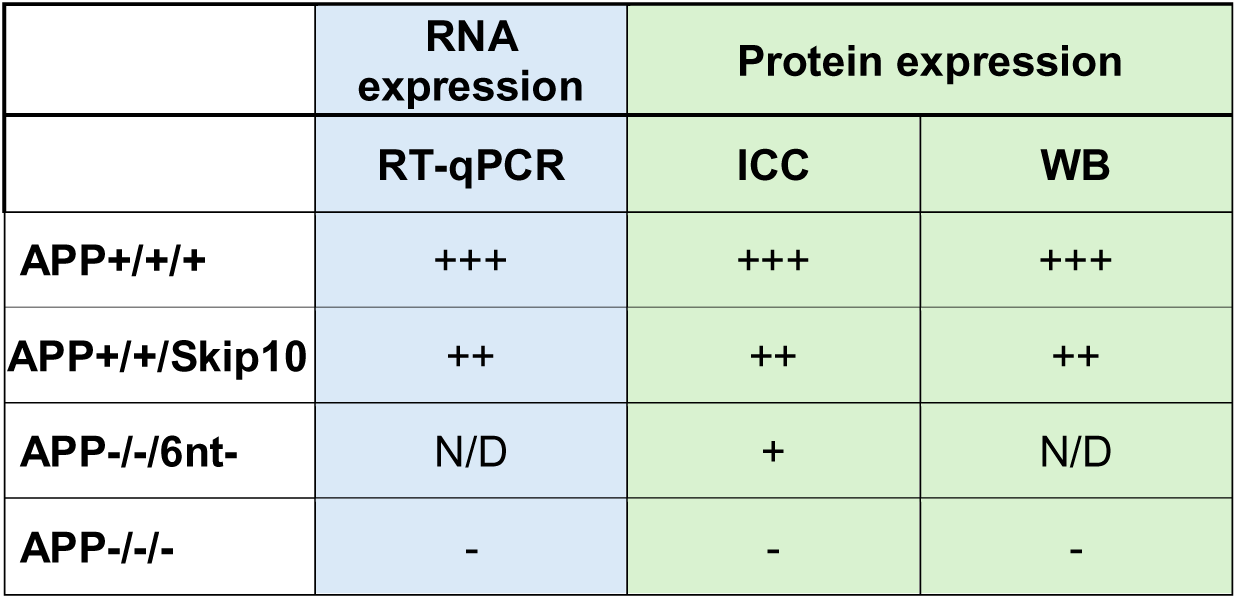
APP expression analysis. Relative level of APP expression using different methods. Immunocytochemistry (ICC) and western blot (WB) analyses were performed on undifferentiated iPS cells. RT-qPCR was done using 92-day organoids grown using each clone. N/D: Not done. Relative expression levels: high (+++), reduced (++) (+), absent (-). (N/D) Not determined.

|  | RNA expression | Protein expression |  |
| --- | --- | --- | --- |
|  | RT-qPCR | ICC | WB |
| **APP+ / + / +** | +++ | +++ | +++ |
| **APP+ / + / Skip10** | ++ | ++ | ++ |
| **APP- / - / 6nt-** | N/D | + | N/D |
| **APP- / - / -** | - | - | - |

*Note: Numerous other APP mutant lines were generated, and this information and cell lines are available upon request*.

### Reduced Terminal Differentiation of Human NPCs to Cortical Neurons in APP disrupted DS-iPSC Cell Lines

To investigate the effect of different APP perturbations on the capacity of DS NPCs to differentiate into cortical neurons, we assessed our panel of isogenic APP-depleted lines using previously established protocols for monolayer differentiation, typically over a 28 day time-course. Using this procedure, trisomic NPCs generally begin to terminally differentiate after day 20 (Czermiński & Lawrence, 2020) **(Fig. S4).** Cultures of the different lines were plated at the same density, grown and differentiated in parallel under identical conditions. The fraction of cells that had terminally differentiated was quantified by calculating the ratio of TUJ1-positive cells (early neuronal marker) to Total/SOX2-positive Neural progenitor cells (NPCs) at 22, 26, and 28 days **(Fig. 2)**. This ratio reflects the balance between cells that had differentiated to neurons or were still undifferentiated progenitors.

The APP-/-/- knock out line most clearly addresses if APP (at some level) is required for the normal progression of this early step in neural differentiation. As evident from **Figure 2**, lack of APP causes a sharp drop in the fraction of cells that had terminally differentiated to neurons at a given time point. While the difference is smaller at day 22, it becomes more pronounced at days 26 and 28, with APP+/+/+ neurons reaching a ratio of 0.63 compared to 0.32 in APP-/-/- cells (day 26) and 0.57 versus 0.31 (day 28).

It was initially unexpected that the APP+/+/Skip10 also showed a marked reduction in neuronal differentiation (e.g. 0.63 vs 0.36 at day 26, and 0.57 vs 0.35 at day 28), as shown in **Figure 2**, despite retaining two intact APP alleles and detectable APP protein. However, sequencing and expression analyses revealed that the donor-inserted exon 10 did not result in non-sense mediated mRNA decay as we initially assumed. Rather, this allele produces an in-frame exon-10-skip transcript encoding a shorter APP protein lacking a substantial segment of the E2 domain. The E2 domain is a structurally important part of APP protein that is involved in many interactions, and the APP protein homodimerizes as well, as further detailed in the Discussion. Hence, the APP protein lacking exon 10 will predictably impact overall protein structure and thus interaction with other proteins or with APP **(see supplemental Figure 3).** Thus, the differentiation defect in this line cannot be attributed to simply reduction of APP dosage as it could very likely be due to interference of the mutant E2-truncated protein with the normal APP or interacting proteins. Other evidence described below further indicates the likely dominant negative effects of the truncated APP protein.

The APP-/-/6nt- cell line also shows impaired differentiation **(Fig. 2)**, but this line differs from APP+/+/Skip10 in that it has more sharply reduced APP dosage, from 3 APP to only one expressed APP allele, and the one APP allele has a very small 6nt in-frame deletion (in exon 9) at the E2 N-terminus. The percent of neurons formed by day 26 and 28 was modestly improved compared to the APP null line, but still strongly reduced compared to the normal trisomic (APP+/+/+) **(Fig. 2)**. While a priori it is possible the 6 nt deletion could impact neurogenesis, results of other analyses below favor that the differentiation defect in this line is most simply explained by reduced APP dosage rather than major disruption of APP structure.

### APP Plays a Role in Human Neuron Structure Required for Normal Neurite Extension

Amyloid precursor protein (APP) has been implicated in neurotrophic functions during neurodevelopment, contributing to neuronal growth and complexity (Coronel et al., 2018; Müller et al., 2017; Sosa et al., 2013, 2017). Since earlier evidence in mice indicated APP plays a role as a non-canonical adhesion protein in the growth cone, here we investigate the participation of APP in neurite outgrowth in human cortical neurons differentiated from our isogenic APP mutant trisomic iPSC lines, to evaluate if APP expression or structural integrity impact early neuronal morphology.

As shown in **Figure 3**, there is a marked deficit in neurite extension for the full APP knock-out line. After 26 days of differentiation, Tuj1-positive neurons from DS-iPSCs (APP+/+/+) have an average neurite length of 191.79 µm whereas (APP-/-/-) DS-iPSC line had a neurite length of 109.52 µm **(Fig. 3)**. A similar marked reduction is seen again at 28 days, when (APP+/+/+) neurons had an average neurite length of 239.48 µm, which is significantly and strongly reduced in the (APP-/-/-) neurons (150.87 µm) **(Fig. 3)**. These results indicate that APP protein plays a role that is required for the proper structural maturation of neurons.

**Figure 3.**
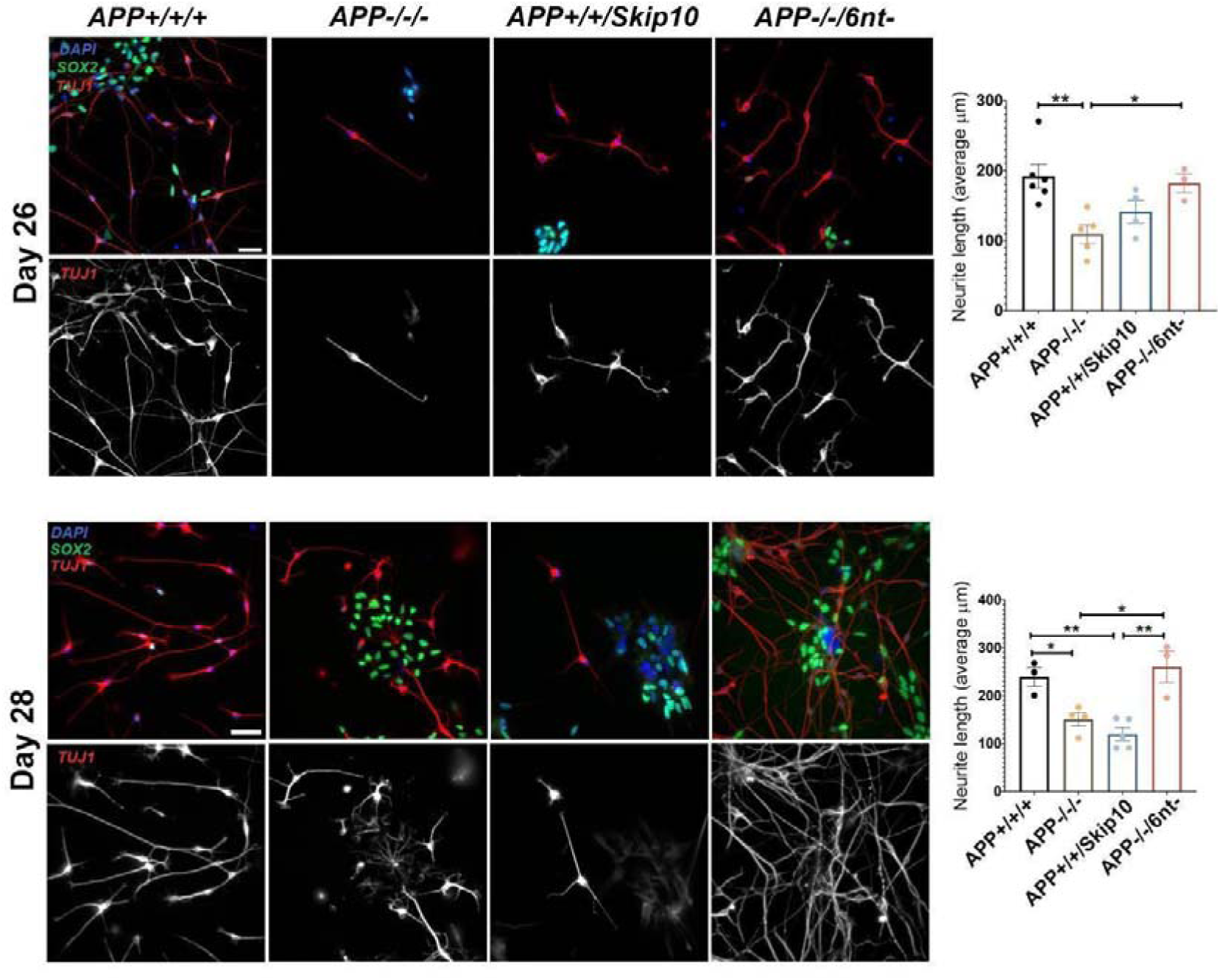
APP depletion/mutation impairs neurite outgrowth in DS-iPSC–derived neurons. Representative immunofluorescence images stained for SOX2 (neural progenitor cells, green) and TUJ1 (neurons, red) of parental DS-iPSCs (APP+/+/+) and isogenic APP depleted lines (APP-/-/-, APP-/-/6nt- and APP+/+/Skip10) after neural differentiation for 26 and 28 days. DAPI (blue) stain nuclei. Scale bar, 50 μm. Measurement of TUJ1+ neurites show neuron derived from APP-depleted lines APP-/-/- and APP+/+/Skip10 exhibit reduced neurite outgrowth compared to APP+/+/+ controls and the APP-/-/6nt- cell line. These results indicate that the short 6 nt mutation present in the APP-/-/6nt- cell line seems to favor neurite outgrowth, while the APP+/+/Skip10 cell line, which has an allele which affects the E2 extracellular domain of APP (which has been shown to be necessary for neurite extension) may have a dominant negative effect on neurite outgrowth.

Neurite length was also substantially reduced for the APP+/+/Skip10 neurons, seen in both Day 26 and Day 28 comparisons. By day 28 the average neurite length was 119.58 for this mutant APP line, much less than 239.48 seen for the APP+++ line. It is perhaps not surprising that the highly abnormal mutant APP protein (lacking all of exon 10) disrupts the structural role of APP required for normal neuron structure. The mutant APP likely also interacts and reduces functionality of the normal APP alleles.

Interestingly, neurons from the APP-/-/6nt- line were not significantly different from the line producing normal APP protein (APP+/+/+) **(Fig. 3).** The average neurite lengths of 182.05 µm at Day 26 and 260.63 at Day 28 compare well to the control cells, but are much longer than those of the APP -/-/- or APP+/+/Skip10 (119.58 µm) neurons. These results showing that the APP-/-/6nt- cells support normal neurite structure indicate that the small in-frame indel has minimal effect on this aspect of APP’s structural role.

Collectively these results indicate that APP is required for proper structural maturation of human neurons. Neurite length is also substantially reduced in APP+/+/Skip10, and at day 28 this phenotype is much closer to APP-/-/- than to APP+/+/+ despite the presence of two intact APP alleles and measurable APP protein.

Because APP+/+/Skip10 expresses both wild-type APP and an in-frame exon-10-skip protein that lacks much of the E2 heparin-binding region, the strong neurite outgrowth defect supports a dominant-negative effect of the mutant protein on the structural functions of wild-type APP. In contrast, APP-/-/6nt- neurons show neurite lengths comparable to APP+/+/+, supporting the interpretation that the 6-nt deletion preserves most E2-dependent neurite-promoting function.

All results together strongly support a structural role of APP in neuronal morphogenesis, which does not tolerate major disruption of E2 domain structure. In accordance with these findings, previous studies suggested that the E2 and extracellular domain of APP is important for cell growth and neurite extension (Müller et al., 2017; Sosa et al., 2017). This region has been implicated in APP dimerization and binding with other proteins, modulating its traffic, recycling, and processing (Eggert et al., 2009; Gustafsen et al., 2013; Ho & Südhof, 2004; Hoe et al., 2009; Müller et al., 2017).

In striking contrast to the exon 10 deletion, in cells in which the only APP had a 6 nt indel (in exon 9) there was no apparent impact on the structural contribution of APP to neurite extension. Hence, this further suggested to us that the decreased differentiation of NPCs to neurons in this line likely reflects the impact of monosomy for APP in trisomic cells. Given that Notch is a well-established regulator of the NPC cycling (Hitoshi et al., 2002; Louvi & Artavanis-Tsakonas, 2006; Ohtsuka, 1999), we next address whether APP dosage may impact NPC differentiation via an effect on Notch, as we further consider in the next section.

### γ-Secretase Inhibition Improves the Impaired Terminal Differentiation of Trisomic Cells Depleted of APP

Since neurite architecture is relatively preserved in APP-/-/6nt- neurons, yet this line exhibits clear reduction in neural progenitor differentiation, these impacts of APP perturbations may involve mechanistically distinct processes. Neurite outgrowth appears primarily dependent on the structural integrity of the APP and its extracellular domain, but we hypothesized that depletion of APP in the null and monosomic lines (APP-/-/6nt-) may underlie the prolonged cycling of neural progenitors (and thus reduction in neuron formation). Specifically, this early neurodevelopmental step may be impacted indirectly by a dosage-dependent effect of APP on maintaining balance with signaling processes. If reduced neurogenesis seen in APP null cells reflects a direct function of APP required for full neurogenic potential, then we would not expect this to be overcome by gamma-secretase inhibition, which is known to trigger neurogenesis from NSCs. Hence, we tested the effects of gamma-secretase inhibition on the APP null line. We also tested the APP-/-/6nt- line, since above results indicate the 6nt deletion does not have a strong or dominant-negative effect on APP function in neuron structure.

This experiment also provides insight into the potential interplay of APP dosage and Notch signaling. Notch signaling drives the cycling of neural progenitor cells, and this requires Notch cleavage by gamma-secretase. For several reasons, there may be complex interplay between APP and Notch-1 trans-membrane receptors, which are primary targets for cleavage by γ-secretase, and thus may compete or otherwise interact (Berezovska et al., 2001; Zhang et al., 2000). Upon γ-secretase cleavage, APP generates the APP intracellular domain (AICD), which has been implicated in transcriptional regulation during neuronal differentiation (Beckett et al., 2012; Coronel et al., 2018). Similarly, γ-secretase-mediated cleavage of Notch releases the Notch intracellular domain (NICD), which translocates to the nucleus and promotes transcription of target genes such as *HES1* and *HES5*, key regulators of progenitor maintenance and inhibition of neuronal differentiation (Hitoshi et al., 2002; Ohtsuka, 1999).

We hypothesized that absent or reduced APP dosage could indirectly affect Notch signaling. Hence, we investigated whether impaired neuronal differentiation in APP-depleted DS-NPCs (APP-/-/-, APP-/-/6nt-) might be mitigated by altered γ-secretase dependent Notch signaling. Alternatively, if APP was required to play a direct role necessary to promote normal NPC differentiation, then gamma-secretase inhibition would not be expected to overcome lack of APP. To test this, we pharmacologically inhibited γ-secretase using Compound E (100 nM) (Beher et al., 2001; Seiffert et al., 2000) during neuronal differentiation. Compound E was added every two days following established differentiation protocols, and neuronal output was quantified at 28 days in vitro. Differentiation efficiency was assessed as the ratio of TUJ1-positive neurons to total cells.

Importantly, on day 28, γ-secretase inhibition significantly increased the TUJ1-positive/total cell ratio in APP-/-/- cultures (0.52) compared with vehicle treated APP-/-/- controls (0.30) **(Fig. 4)**. Hence, the lower levels of neurogenesis in cells which lack any APP can be overcome by inhibition of Notch signaling. A similar effect was observed in APP-/-/6nt- cultures, in which the ratio of neurons increased from 0.38 (vehicle treated) to 0.53 following Compound E treatment. These values approached those observed in untreated APP+/+/+ cultures, indicating a rescue of the differentiation deficit. While we cannot rule out that there may be some other aspect of APP-dependent regulation, results suggest that the requirement for APP may be largely through indirect impacts on Notch signaling and NPC cycling.

**Figure 4.**
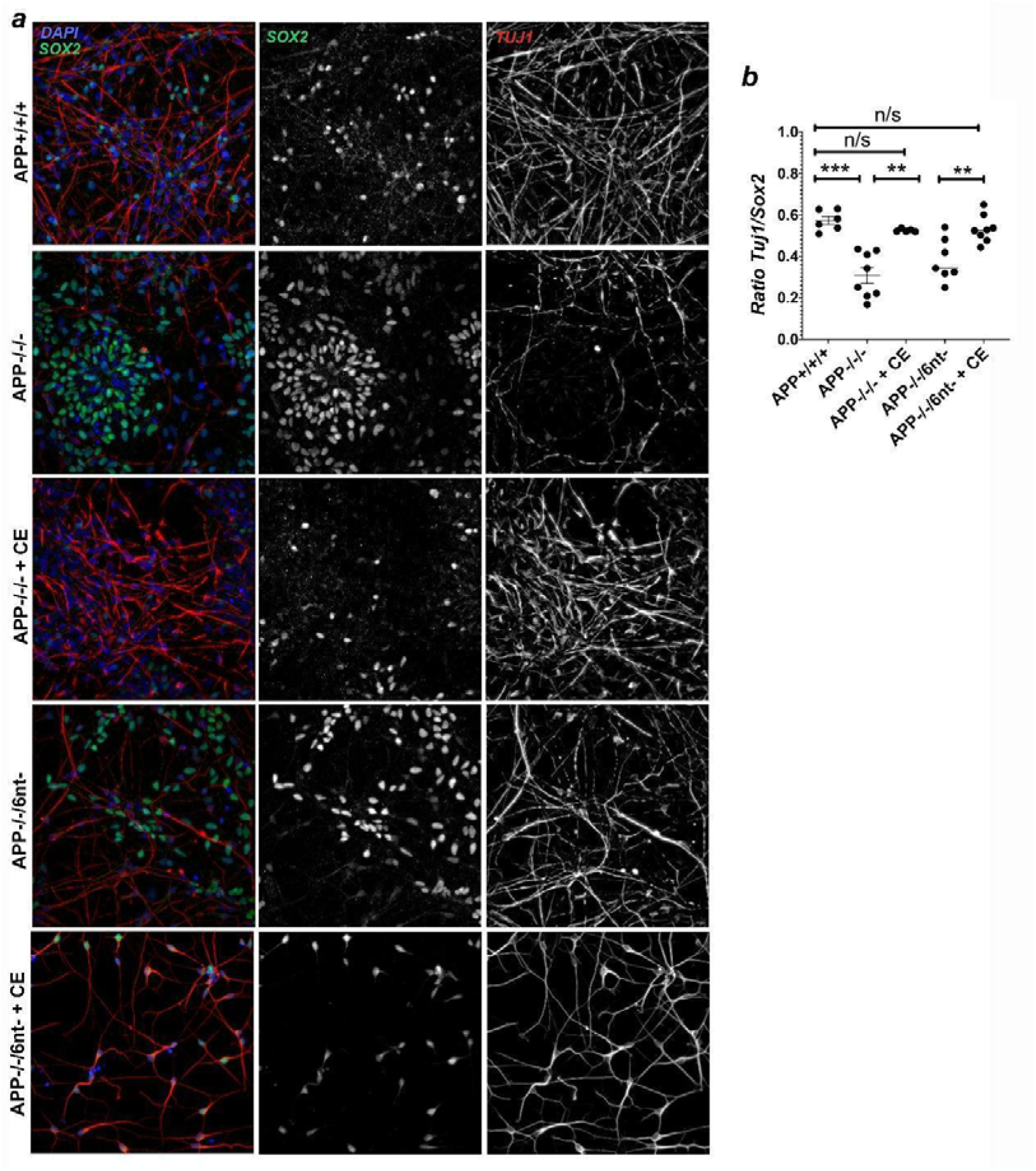
γ-Secretase inhibition restores neuronal differentiation in APP-depleted DS-iPSC lines. (a) Representative images of parental DS-iPSC lines (APP+/+/+) and their isogenic APP-depleted variants APP-/-/-, APP-/-/6nt-, either untreated or treated with Compound E (100 nM), stained for SOX2 (neural progenitor marker) and TUJ1 (neuronal marker) at day 28 of differentiation. DAPI (blue) marks nuclei. (b) Ratio of TUJ1⁺/SOX2⁺ cells are significantly increased in Compound E–treated APP-/-/-, APP-/-/6nt- lines compared to untreated conditions, indicating enhanced neuronal differentiation.

As considered in the Discussion, there is evidence that other chromosome 21 genes may influence Notch signaling in the context of Trisomy 21 (Moon & Lawrence, 2022), but these results provide evidence that the interplay of APP and Notch signaling is a key aspect.

## DISCUSSION

This study contributes to the large body of work investigating the biology of APP, specifically with a focus here on the impact of perturbing APP in the context of Trisomy 21. Using a unique panel of DS-iPSC depleted and mutated APP cell lines, we provide evidence that APP exerts dual and mechanistically distinct functions during early DS neurodevelopment: an indirect dosage-dependent role in neural progenitor fate regulation and a direct structure-dependent role in neuron architecture and neurite extension.

We first demonstrate an early impairment in the differentiation of NPC to neurons with an increase in Sox2 positive NSCs in depleted APP DS-iPS cell lines. Notably, the Notch signaling pathway also supports the undifferentiated state of NPCs (Hitoshi et al., 2002; Louvi & Artavanis-Tsakonas, 2006; Ohtsuka, 1999), suggesting an APP mediated dysregulation of Notch signaling may be involved. The fact that APP-/-/6nt- also has an effect on NSC cycling, but not on neurite structure, further suggests a mechanism whereby APP dosage reduction indirectly impacts NPC differentiation to neurons.

Molecular interactions between APP and Notch receptors have been previously described (Chen et al., 2006; Oh et al., 2005). The interaction between APP and Notch receptors can occur in cis, where both transmembrane proteins reside within the same cellular compartment and compete for the pool of γ-secretase. Such competition between Notch and APP has been demonstrated (Berezovska et al., 2001; Lleó et al., 2003; Zhang et al., 2000) as well as between Notch and Delta (LaVoie & Selkoe, 2003). To investigate the involvement of Notch signaling in the reduction of NSC differentiation to neurons observed in depleted APP DS-iPSC cell lines, we pharmacologically inhibited γ-secretase activity using compound E. This intervention led to a notable rescue of neuronal differentiation propensity; hence the lack of APP can be overcome by a drug which impacts Notch cleavage and signaling. This is consistent with the depletion of APP impacting Notch signaling homeostasis and thus the balance between NPC maintenance and early neuronal differentiation.

Results here provide a complementary perspective to the accelerated differentiation reported in euploid cells fully depleted for APP in an in-depth study by (Shabani et al., 2023). Importantly, both studies agree that lack of APP impacts early steps in neurogenesis. Although there are differences in the directional impact of APP loss on the balance of NPC and neuron differentiation, such differences likely reflect that the developmental role of APP is context dependent. Foremost, our cells carry a trisomy 21 background where baseline Notch signaling is already elevated by over-expression of chromosome 21 genes, as shown previously (Czermiński & Lawrence, 2020; Moon & Lawrence, 2022), and our other recent findings strongly implicate another chr21 gene(s) as driving increased Notch signaling (Larsen et al. in prep). While our analyses emphasize a prospective effect of APP on Notch cleavage and signaling, supported by the Compound E rescue experiment, there is a complex network of signaling pathways.

For example, in euploid cells with balanced baseline Notch levels, the impact on WNT/AP-1 signaling may be prominent. Second, the two studies capture distinct temporal windows of neuron development: (Shabani et al., 2023)analyzed early progenitor fate decisions (days 0-7), while we examined terminal differentiation and structural maturation at a later stage (days 22-28). Despite these differences in experimental design and timing, both studies show that lack of APP impacts this early step in neurogenesis. In accordance with these early developmental impacts, we note a rare clinical case involving a homozygous nonsense mutation in APP (resulting in a truncated protein), led to profound neurodevelopmental abnormalities (Klein et al., 2016)

The second effect that we found in the APP mutant lines is a direct structure-dependent role for APP in neuron architecture and neurite extension. Our findings are consistent with previous reports describing APP as a type I transmembrane glycoprotein with structural features characteristic of a cell adhesion molecule, essential for promoting neurite outgrowth and cell migration (Coronel et al., 2018; Gabriele et al., 2022; Müller et al., 2017; Sosa et al., 2017). It is known that the large extracellular N-terminal region of APP, encompassing the E1 and E2 domains, plays a crucial role in dimerization, mediating cell adhesion and neurite outgrowth functions in mouse neurons (Müller et al., 2017; Sosa et al., 2017). The E2 domain constitutes a large structure within the extracellular ectodomain, which is encoded, in part, by Exons 9 and 10 **(Fig. S3)**. The APP-/-/6nt- cells harbor an in-frame micro-deletion within Exon 9, changing the conserved AVD sequence to a single D residue which likely preserves E2 structure and retains trophic function. In contrast, APP+/+/Skip10 large deletion significantly impacts one of the heparin-binding sequences in region required to orchestrate cis and trans homodimerization, and impacting interactions with cell-surface receptors (e.g. LRP1 and SorLA) to guide retrograde trafficking (Eggert et al., 2009; Gustafsen et al., 2013; Müller et al., 2017; Sosa et al., 2017).

The different cellular phenotypes associated with different mutants can provide insights into APP biology. Although *APP+/+/Skip10* neurons express three *APP* alleles, deletion of a substantial portion of the E2 domain in one allele apparently acts in a dominant-negative manner, disrupting the adhesive, dimeric, and trafficking mechanisms essential for neurite outgrowth (Dahms et al., 2012; Jin et al., 1994; Multhaup et al., 1994; Reinhard et al., 2005). In contrast, *APP-/-/6nt-* neurons, which express a single allele missing only two amino acids, exhibit neurite extensions comparable to wild-type *APP+/+/+* neurons, indicating preservation of overall E2 domain structure sufficient to sustain adhesive capacities required for normal neurite elongation.

Together, these findings position APP as a dosage-sensitive regulator of progenitor maintenance within the trisomic context, while simultaneously acting as a structural determinant of neuronal morphogenesis through its extracellular domain. This functional separation refines our understanding of APP biology in the context of trisomy 21 and also raises important considerations that therapeutic interventions for APP must be carefully evaluated to avoid unintended disruptions of signaling homeostasis.

## KEY RESOURCES TABLE

### Antibodies

| REAGENT or RESOURCE | SOURCE | IDENTIFIER |
| --- | --- | --- |
| Anti-Sox2 | Millipore | Cat# AB5603;<br>RRID:AB_2286686 |
| Anti-TUBB3 | BioLegend | Cat# 801202;<br>RRID:AB_10063408; Tuj1 |
| Anti-APP | Sigma-Aldrich | Cat# MAB348 |

### Chemicals, Peptides, and Recombinant Proteins

|  | SOURCE | IDENTIFIER |
| --- | --- | --- |
| DMH1 | Tocris Bioscience | Cat# 4126 |
| SB 431542 | Tocris Bioscience | Cat# 1614 |
| Recombinant Human GDNF | Preprotech | Cat# 4050-10 |
| Recombinant Human/Murine/Rat BDNF | Preprotech | Cat# 4050-02 |
| Y-27632 | Tocris Bioscience | Cat# 1254 |
| Vitronectin (VTN-N) Recombinant Human Protein, Truncated | Thermo Fisher Scientific | Cat# A14700 |
| L-Ascorbic Acid | Sigma Aldrich | Cat# A4544 |
| Adenosine 3<sup>0</sup>,5<sup>0</sup>-cyclic monophosphate sodium salt monohydrate (cyclic AMP) | Sigma Aldrich | Cat# A6885 |
| Compound E | Millipore | Cat# 565790 |
| Essential 8 Medium | Thermo Fisher Scientific | Cat# A1517001 |
| B-27 Supplement (50X), | Thermo Fisher Scientific | Cat# 12587010 |
| **minus vitamin A** |  |  |
| **N-2 Supplement (100X)** | Thermo Fisher Scientific | Cat# 17502048 |
| **Neurobasal Medium** | Thermo Fisher Scientific | Cat# 21103049 |
| **Knockout DMEM/F-12** | Thermo Fisher Scientific | Cat# 12660012 |
| **UltraPure 0.5M EDTA, pH 8.0** | Thermo Fisher Scientific | Cat# 15575020 |
| **Laminin** | Sigma Aldrich | Cat# L2020-1MG |

### RNA extraction and RT-qPCR reagent / Kit

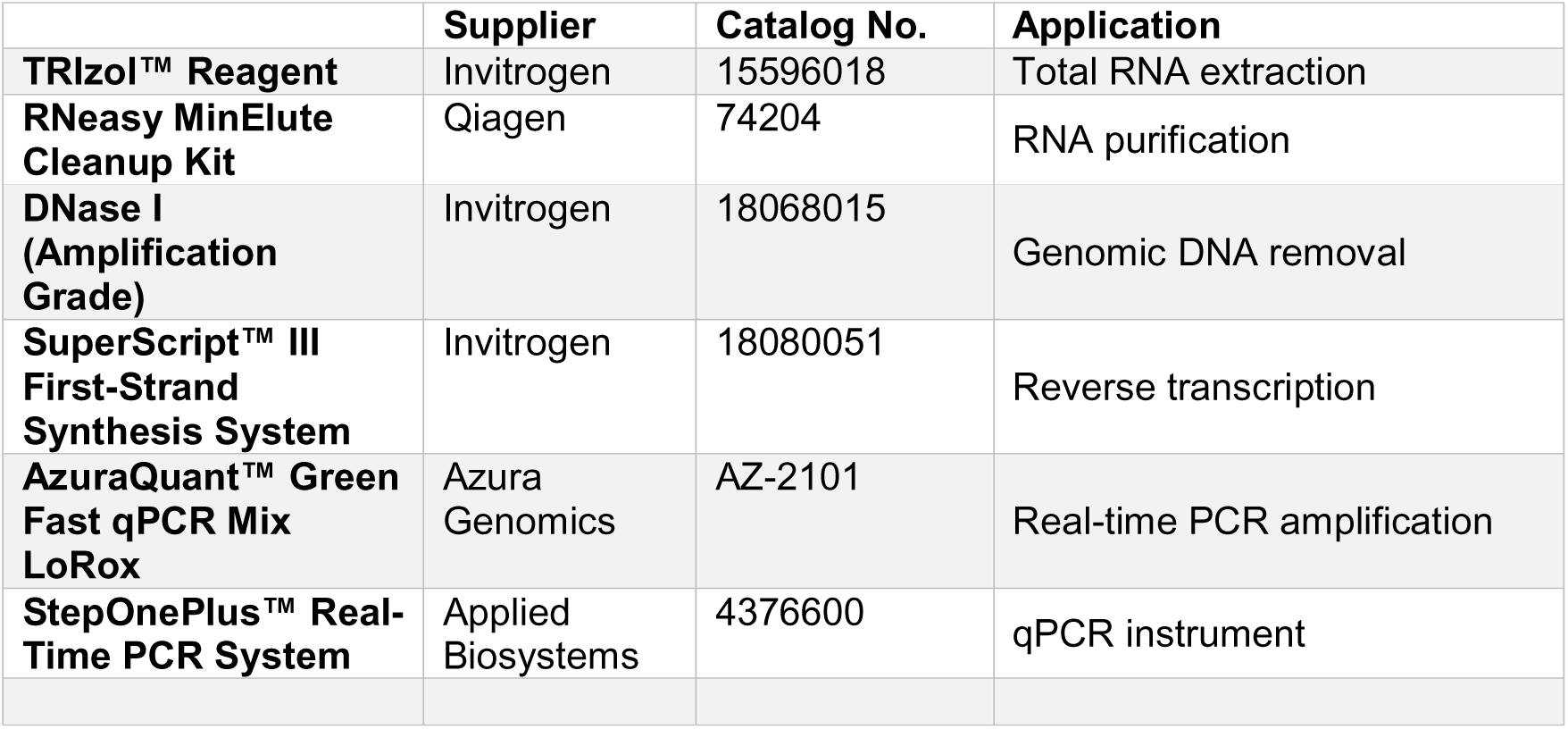

|  | **Supplier** | **Catalog No.** | **Application** |
| --- | --- | --- | --- |
| **TRIzol™ Reagent** | Invitrogen | 15596018 | Total RNA extraction |
| **RNeasy MinElute Cleanup Kit** | Qiagen | 74204 | RNA purification |
| **DNase I (Amplification Grade)** | Invitrogen | 18068015 | Genomic DNA removal |
| **SuperScript™ III First-Strand Synthesis System** | Invitrogen | 18080051 | Reverse transcription |
| **AzuraQuant™ Green Fast qPCR Mix LoRox** | Azura Genomics | AZ-2101 | Real-time PCR amplification |
| **StepOnePlus™ Real-Time PCR System** | Applied Biosystems | 4376600 | qPCR instrument |

### sgRNA and oligo sequence for CRISPR/Cas9

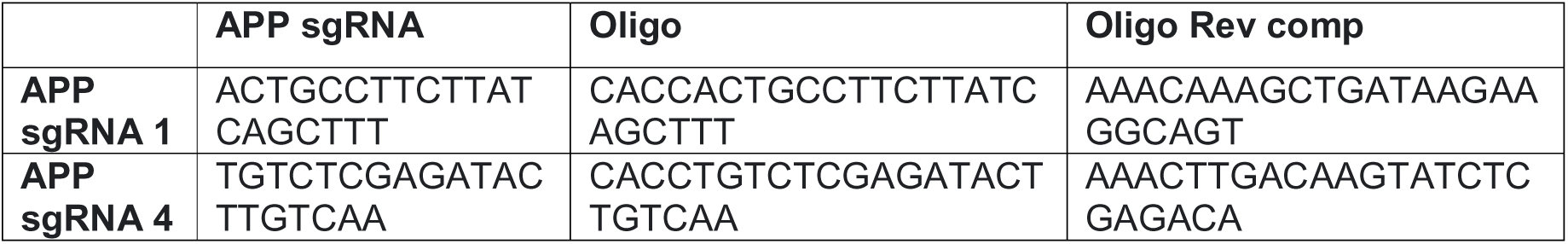

### Primer sequence for RT qPCR

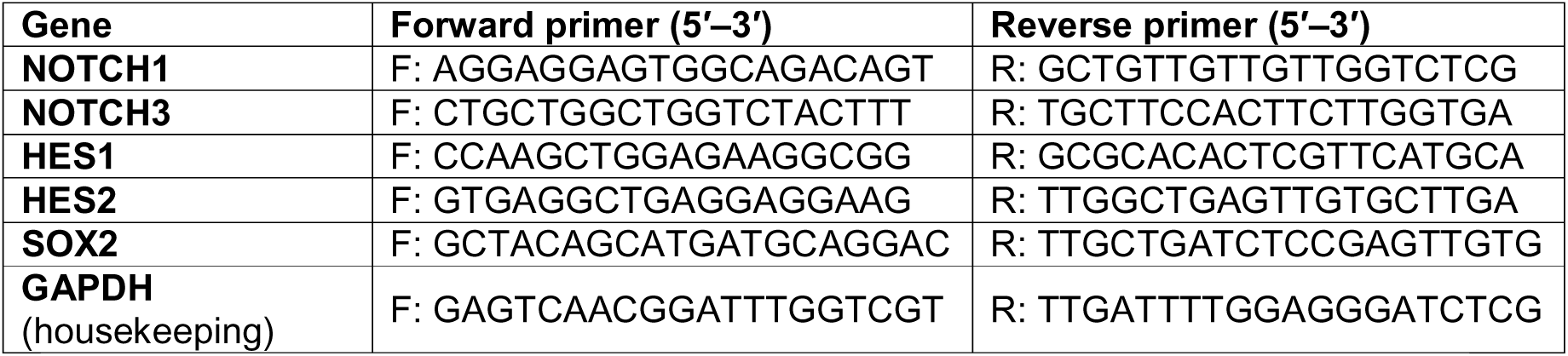

## EXPERIMENTAL MODEL AND SUBJECT DETAILS

### Human Cells

The initial DS iPSC parental line (DS1-iPS4) was provided by G.Q. Daley (Park et al., 2008) and is derived from a 1-year-old male with Down syndrome. The isogenic APP depleted subclones were derived from the DS iPSC parental line editing by CRISPR/Cas9. iPSCs were maintained on vitronectin-coated plates with Essential 8 medium (ThermoFisher) at 37C with 20% O2 and 5% CO2. Cells were tested periodically for mycoplasma and passed every 3-4 days with 0.5mM EDTA.

### DS-IPSCs cell line editing

In order to reduce the copy numbers/expression of APP on trisomy/Parental ***DS-iPSCs*** cell lines (APP+/+/+), we used CRISPR/Cas9 genomic editing technology to create dsDNA break. Thus, we used CRISPR/Cas9 together with a donor construct with selection cassette and perform homolog recombination to produce a truncated 5’ end mRNA susceptible to be degraded by exoribonucleases **(Fig. 1a).** For that It was used a pCRISPR/Cas-GFP with sgRNA cloning site (pSpCas9(BB)-2A-GFP-APP). The guide RNA, sgRNA 1 and 2 correspond to EXON 10 and sgRNA3 and 4 for EXON 9 of APP. We first transfected the cell lines with the plasmid for CRISPR/Cas9 and then after checking the transfection, we performed a co-transfection with the plasmid for CRISPR/Cas9 and the donor DNA to cells (pRedPuro TK Exon 10 HA Assembled) and (pRedPuro TK Exon 9 HA Assembled). DNA ratio was 1:1 sgRNA + pRedPuro TK Exon 10 HA Assembled and sgRNA 4 + pRedPuro TK Exon 9 HA Assembled.

After transfection cells were treated with 0.3 μg/ml puromycin (Puro) to select for the pRedPuro insert. The Puro selection was performed: 2 days after transfection -0.2 ug/ml for 3 days. We let cells recover and start again with 0.2 ug/ml, until cells look stable, then 3 ug/ml, then 4 ug/ml for 4 weeks. After a month 2 mM ganciclovir (GCV) for a week to eliminate cells with pRedPuroTK random integration.

## METHOD DETAILS

### Neural Differentiation

Neural differentiations were performed as previously described (Cao et al., 2017; Chambers et al., 2009) with some modifications. Briefly, iPSCs were dissociated into single cells and plated at a density of 50,000 cells/well in a vitronectin-coated 24-well plate with 10mM of the ROCK inhibitor Y-27632 (Tocris Bioscience). The next day, media was changed to Neural differentiation media (NDM) consisting of 50% DMEM/F12, 50% Neurobasal, 0.5X Glutamax, 1X N-2 supplement, 1X penicillin/streptomycin (all from ThermoFisher), and supplemented with 2uM DMH1 and SB431542 (both from Tocris Bioscience). After 14 days, cells were broken into clumps after EDTA treatment and cultured in suspension for 7 days in NDM. On diff21 or diff28, neurospheres were dissociated into single cells with StemPro Accutase (ThermoFisher) and plated onto coverslips (Electron Microscopy Sciences) coated with Laminin (Sigma-Aldrich, Cat#L2020-1MG) at 1ug/ml and a density of 25,000-50,000 cells/coverslip and fed every 2-3 days with Neuron media consisting of Neurobasal, 1X N-2, 0.5X B-27 without vitamin A, 1X penicillin/streptomycin, 1X Glutamax (ThermoFisher), 0.3% Glucose, 10ng/ml GDNF (Peprotech), 10ng/ml BDNF (Peprotech), 10ng/ml ascorbic acid (Sigma-Aldrich), and 1mM cyclic AMP (Sigma-Aldrich). In cultures where NPC differentiation was studied, compound E (EMD Millipore) was added for 2 days at diff23 or diff25 at a concentration of 100nM.

### Cell Fixation, RNA FISH, and Immunofluorescence

For iPSC and monolayer neural culture, cell fixation with 4% paraformaldehyde (PFA) was performed as previously described (Byron et al., 2013). RNA FISH and IF were performed as previously described (Byron et al., 2013). For RNA FISH and combined RNA FISH with IF in iPSCs culture, detergent extraction was performed prior to fixation. For IF alone, fixation was performed prior to detergent extraction. The APP probe is a BAC from BACPAC resources (RP11-910G8). The primary antibodies used in this study are provided in the Key Resources. The conjugated secondary antibodies used in this study were Alexa Fluor 488, 594, and 647.

### RNA extraction and RT-qPCR

Total RNA was extracted from cultured cells using TRIzol™ reagent (Invitrogen) according to the manufacturer’s protocol. RNA was further purified with RNeasy MinElute columns (Qiagen) and treated with DNase I (Amplification Grade, Invitrogen) to remove genomic DNA contamination. Reverse transcription was performed with the SuperScript™ III First-Strand Synthesis System (Invitrogen) using oligo (dT) 20 primers, following the supplier’s instructions. Quantitative PCR was performed using a StepOnePlus™ Real-Time PCR System (Applied Biosystems). Amplifications were carried out with AzuraQuant™ Green Fast qPCR Mix LoRox (Azura Genomics, USA) in a total reaction volume of 20 µL containing 10 µL of 2× master mix, 0.8 µL each of forward and reverse primers (10 µM, final concentration 400 nM), template cDNA (<100 ng), and nuclease-free water. Primer sequences for target genes (NOTCH1, NOTCH3, HES1, HES2, and SOX2) and the housekeeping gene GAPDH were designed and synthesized by Integrated DNA Technologies (USA). Cycling conditions consisted of an initial enzyme activation step at 95 °C for 2 min, followed by 40 cycles of denaturation at 95 °C for 5 s and annealing/extension at 60 °C for 20 s. Melting curve analysis was performed at the end of each run to confirm amplification specificity. Each reaction was carried out in triplicate. Relative expression levels were calculated using the 2ΔΔCt method with GAPDH as an internal control.

### Microscopy

Cells were visualized using a Zeiss Axio Observer 7, equipped with Chroma multi-bandpass dichroic and emission filter sets (Brattleboro, VT), with a Flash 4.0 LTCMOS camera (Hamamatsu). Brightness and contrast were corrected in Fiji to best represent what was observed by eye.

### Quantification and Statistical analysis

#### Microscopy Quantification

For scoring of neuron versus NPC cell type, TUJ1+ cells were counted as neurons, SOX2 cells were counted as NPC, and SOX2-/TUJ1 cells were not counted. For these experiments, 6 random low-power fields from different cell lines differentiated 1-2 times were examined for each condition. Neurite length of TUJ1+ neurons cells was measured using in Fiji.

#### Statistics

All statistical analyses were performed, and graphs generated using GraphPad Prism 8. Data are shown as mean ± SEM unless otherwise stated. In general, all comparisons for groups of three or more were analyzed by one way ANOVA followed by a Tukey’s multiple comparison tests. Pairwise sample comparisons were performed using Student’s t-test.

## Acknowledgements

Work done by LJS on this study was performed at UMass Chan Medical School. We thank Eric Larsen for helpful assistance and discussions. We thank Meg Byron, Khushali Gupta, and Yuanchun Jing for general lab and technical assistance and for their support as needed.

## Data availability statement

The raw data supporting the conclusions of this article will be made available by the authors, without undue reservation.

## Ethics statement

The study was conducted in accordance with the local legislation and institutional requirements.

## Author contributions

MV designed and performed genetic engineering experiments and did molecular analyses to characterize specific mutations in all iPSC lines, LJS planned and performed analyses of neural differentiation in all lines, interpreted results and drafted the manuscript, KS contributed to data analysis and manuscript preparation, and JBL initiated the project, contributed to interpreting results, and edited the manuscript . All authors contributed to editing and approved the final submitted manuscript.

## Funding

The author(s) declare that financial support was received for the research, authorship, and/or publication of this article. This research was supported by funding from NIH grant R01HD091357 to JBL and Alz. Assoc. Grant #24AARG-D-1198629 to LJS. MV received additional support from NIH grant F32AG056131.

## Conflict of interest

The authors declare that the research was conducted in the absence of any commercial or financial relationships that could be construed as a potential conflict of interest.

## Notes

### Competing Interest Statement

The authors have declared no competing interest.

## REFERENCES

1. Beckett, C., Nalivaeva, N. N., Belyaev, N. D., & Turner, A. J. (2012). Nuclear signalling by membrane protein intracellular domains: The AICD enigma. *Cellular Signalling*, Feb;24(2):402–409. 10.1016/j.cellsig.2011.10.007

2. Beher, D., Wrigley, J. D. J., Nadin, A., Evin, G., Masters, C. L., Harrison, T., Castro, J. L., & Shearman, M. S. (2001). Pharmacological knock-down of the presenilin 1 heterodimer by a novel γ-secretase inhibitor: Implications for presenilin biology. Journal of Biological Chemistry, 276(48). 10.1074/jbc.M103075200

3. Berezovska, O., Jack, C., Deng, A., Gastineau, N., Rebeck, G. W., & Hyman, B. T. (2001). Notch1 and Amyloid Precursor Protein Are Competitive Substrates for Presenilin1-dependent γ-Secretase Cleavage. Journal of Biological Chemistry, 276(32). 10.1074/jbc.M008268200

4. Bolós, M., Hu, Y., Young, K. M., Foa, L., & Small, D. H. (2014). Neurogenin 2 mediates amyloid-β precursor protein-stimulated neurogenesis. Journal of Biological Chemistry, 289(45). 10.1074/jbc.M114.581918

5. Byron, M., Hall, L. L., & Lawrence, J. B. (2013). A multifaceted FISH approach to study endogenous RNAs and DNAs in native nuclear and cell structures. *Current Protocols in Human Genetics*, Jan;Chapter 4:Unit 4.15. 10.1002/0471142905.hg0415s76

6. Callahan, D. G., Taylor, W. M., Tilearcio, M., Cavanaugh, T., & Selkoe, D. J. (2017). Embryonic mosaic deletion of APP results in displaced Reelin-expressing cells in the cerebral cortex. Developmental Biology, 424(2), 138–146. 10.1016/j.ydbio.2017.03.007

7. Cao, S. Y., Hu, Y., Chen, C., Yuan, F., Xu, M., Li, Q., Fang, K. H., Chen, Y., & Liu, Y. (2017). Enhanced derivation of human pluripotent stem cell-derived cortical glutamatergic neurons by a small molecule. Scientific Reports, 7(1). 10.1038/s41598-017-03519-w

8. Chambers, S. M., Fasano, C. A., Papapetrou, E. P., Tomishima, M., Sadelain, M., & Studer, L. (2009). Highly efficient neural conversion of human ES and iPS cells by dual inhibition of SMAD signaling. Nature Biotechnology, 27(3). 10.1038/nbt.1529

9. Chen, C. Di, Oh, S. Y., Hinman, J. D., & Abraham, C. R. (2006). Visualization of APP dimerization and APP-Notch2 heterodimerization in living cells using bimolecular fluorescence complementation. Journal of Neurochemistry, 97(1). 10.1111/j.1471-4159.2006.03705.x

10. Coronel, R., Bernabeu-Zornoza, A., Palmer, C., Muñiz-Moreno, M., Zambrano, A., Cano, E., & Liste, I. (2018). Role of Amyloid Precursor Protein (APP) and Its Derivatives in the Biology and Cell Fate Specification of Neural Stem Cells. *Molecular Neurobiology*, Sep;55(9):7107–7117. 10.1007/s12035-018-0914-2

11. Czermiński, J. T., & Lawrence, J. B. (2020). Silencing Trisomy 21 with XIST in Neural Stem Cells Promotes Neuronal Differentiation. Developmental Cell, 52(3), 294–308.e3. 10.1016/j.devcel.2019.12.015

12. Dahms, S. O., Könnig, I., Roeser, D., Gührs, K. H., Mayer, M. C., Kaden, D., Multhaup, G., & Than, M. E. (2012). Metal binding dictates conformation and function of the amyloid precursor protein (APP) E2 domain. Journal of Molecular Biology, 416(3). 10.1016/j.jmb.2011.12.057

13. Dunot, J., Ribera, A., Pousinha, P. A., & Marie, H. (2023). Spatiotemporal insights of APP function. Current Opinion in Neurobiology Oct:82:102754. 10.1016/j.conb.2023.102754

14. Eggert, S., Midthune, B., Cottrell, B., & Koo, E. H. (2009). Induced dimerization of the amyloid precursor protein leads to decreased amyloid-β protein production. Journal of Biological Chemistry, 284(42). 10.1074/jbc.M109.038646

15. Fortea, J., Vilaplana, E., Carmona-Iragui, M., Benejam, B., Videla, L., Barroeta, I., … Lleó, A. (2020). Clinical and biomarker changes of Alzheimer’s disease in adults with Down syndrome: a cross-sectional study. The Lancet, 395(10242), 1988–1997. 10.1016/S0140-6736(20)30689-9

16. Gabriele, R. M. C., Abel, E., Fox, N. C., Wray, S., & Arber, C. (2022). Knockdown of Amyloid Precursor Protein: Biological Consequences and Clinical Opportunities. In Frontiers in Neuroscience (Vol. 16). 10.3389/fnins.2022.835645

17. Gupta, K., Czerminski, J. T., & Lawrence, J. B. (2024). Trisomy silencing by XIST: translational prospects and challenges. Human Genetics Mar 9;143(7):843–855. 10.1007/s00439-024-02651-8

18. Gustafsen, C., Glerup, S., Pallesen, L. T., Olsen, D., Andersen, O. M., Nykjær, A., Madsen, P., & Petersen, C. M. (2013). Sortilin and SorLA display distinct roles in processing and trafficking of amyloid precursor protein. Journal of Neuroscience, 33(1), 64–71. 10.1523/jneurosci.2371-12.2013

19. Hitoshi, S., Alexson, T., Tropepe, V., Donoviel, D., Elia, A. J., Nye, J. S., Conlon, R. A., Mak, T. W., Bernstein, A., & Van Der Kooy, D. (2002). Notch pathway molecules are essential for the maintenance, but not the generation, of mammalian neural stem cells. Genes and Development, 16(7). 10.1101/gad.975202

20. Ho, A., & Südhof, T. C. (2004). Binding of F-spondin to amyloid-β precursor protein: A candidate amyloid-β precursor protein ligand that modulates amyloid-β precursor protein cleavage. Proceedings of the National Academy of Sciences USA, 101(8). 10.1073/pnas.0308655100

21. Hoe, H., Lee, K. J., Carney, R. S. E., Lee, J., Lee, J., Howell, B. W., Hyman, B. T., Pak, D. T. S., & Rebeck, G. W. (2009). Interaction of Reelin with APP promotes neurite outgrowth. Journal of Neuroscience, 29(23), 7459–7473. 10.1523/jneurosci.4872-08.2009.

22. Hu, Y., Hung, A. C., Cui, H., Dawkins, E., Bolós, M., Foa, L., Young, K. M., & Small, D. H. (2013). Role of cystatin C in amyloid precursor protein-induced proliferation of neural stem/progenitor cells. Journal of Biological Chemistry, 288(26). 10.1074/jbc.M112.443671

23. Jin, L. W., Ninomiya, H., Roch, J. M., Schubert, D., Masliah, E., Otero, D. A. C., & Saitoh, T. (1994). Peptides containing the RERMS sequence of amyloid β/A4 protein precursor bind cell surface and promote neurite extension. Journal of Neuroscience, 14(9). 10.1523/jneurosci.14-09-05461.1994

24. Klein, S., Goldman, A., Lee, H., Ghahremani, S., Bhakta, V., Nelson, S. F., & Martinez-Agosto, J. A. (2016). Truncating mutations in APP cause a distinct neurological phenotype. Annals of Neurology, 80(3). 10.1002/ana.24727

25. LaVoie, M. J., & Selkoe, D. J. (2003). The Notch Ligands, Jagged and Delta, Are Sequentially Processed by α-Secretase and Presenilin/γ-Secretase and Release Signaling Fragments. Journal of Biological Chemistry, 278(36). 10.1074/jbc.M302659200

26. Lleó, A., Berezovska, O., Ramdya, P., Fukumoto, H., Raju, S., Shah, T., & Hyman, B. T. (2003). Notch1 Competes with the Amyloid Precursor Protein for γ-Secretase and Down-regulates Presenilin-1 Gene Expression. Journal of Biological Chemistry, 278(48). 10.1074/jbc.M308480200

27. Louvi, A., & Artavanis-Tsakonas, S. (2006). Notch signalling in vertebrate neural development. In Nature Reviews Neuroscience Feb;7(2):93-102. 10.1038/nrn1847

28. Moon, J. E., & Lawrence, J. B. (2022). Chromosome silencing in vitro reveals trisomy 21 causes cell-autonomous deficits in angiogenesis and early dysregulation in Notch signaling. Cell Reports, 40(6). 10.1016/j.celrep.2022.111174

29. Müller, U. C., Deller, T., & Korte, M. (2017). Not just amyloid: Physiological functions of the amyloid precursor protein family. In *Nature Reviews Neuroscience*, May;18(5):281–298. 10.1038/nrn.2017.29

30. Multhaup, G., Bush, A. I., Pollwein, P., & Masters, C. L. (1994). Interaction between the zinc(II) and the heparin binding site of the Alzheimer’s disease βA4 amyloid precursor protein (APP). FEBS Letters, 355(2). 10.1016/0014-5793(94)01176-1

31. Oh, S. Y., Ellenstein, A., Di Chen, C., Hinman, J. D., Berg, E. A., Costello, C. E., Yamin, R., Neve, R. L., & Abraham, C. R. (2005). Amyloid precursor protein interacts with notch receptors. Journal of Neuroscience Research, 82(1). 10.1002/jnr.20625

32. Ohtsuka, T. (1999). Hes1 and Hes5 as Notch effectors in mammalian neuronal differentiation. The EMBO Journal, 18(8). 10.1093/emboj/18.8.2196

33. Park, I. H., Arora, N., Huo, H., Maherali, N., Ahfeldt, T., Shimamura, A., Lensch, M. W., Cowan, C., Hochedlinger, K., & Daley, G. Q. (2008). Disease-Specific Induced Pluripotent Stem Cells. Cell, 134(5). 10.1016/j.cell.2008.07.041

34. Peralta Cuasolo, Y. M., Dupraz, S., Unsain, N., Bisbal, M., Quassollo, G., Galiano, M. R., Grassi, D., Quiroga, S., & Sosa, L. J. (2023). The GTPase Rab21 is required for neuronal development and migration in the cerebral cortex. *Journal of Neurochemistry*, Sep;166(5):790–808. 10.1111/jnc.15925

35. Reinhard, C., Hébert, S. S., & De Strooper, B. (2005). The amyloid-β precursor protein: Integrating structure with biological function. *EMBO Journal*, Dec 7;24(23):3996–4006. 10.1038/sj.emboj.7600860

36. Rovelet-lecrux, A., Hannequin, D., Raux, G., Vital, A., Le Meur, N., Laquerrie, A., & Vercelletto, M. (2006). APP locus duplication causes autosomal dominant early-onset Alzheimer disease with cerebral amyloid angiopathy. Nature Genetics 38(1): 24–6. 10.1038/ng1718

37. Seiffert, D., Bradley, J. D., Rominger, C. M., Rominger, D. H., Yang, F., Meredith, J. E., Wang, Q., … Zaczek, R. (2000). Presenilin-1 and -2 are molecular targets for γ-secretase inhibitors. Journal of Biological Chemistry, 275(44). 10.1074/jbc.M005430200

38. Shabani, K., Pigeon, J., Benaissa Touil Zariouh, M., Liu, T., Saffarian, A., Komatsu, J., Liu, E., Danda, N., Becmeur-Lefebvre, M., Limame, R., Bohl, D., Parras, C., & Hassan, B. A. (2023). The temporal balance between self-renewal and differentiation of human neural stem cells requires the amyloid precursor protein. Science Advances Jun 16;9(24). 10.1126/sciadv.add5002

39. Sosa, L. J., Bergman, J., Estrada-Bernal, A., Glorioso, T. J., Kittelson, J. M., & Pfenninger, K. H. (2013). Amyloid Precursor Protein Is an Autonomous Growth Cone Adhesion Molecule Engaged in Contact Guidance. PLoS ONE, 8(5). 10.1371/journal.pone.0064521

40. Sosa, L. J., Cáceres, A., Dupraz, S., Oksdath, M., Quiroga, S., & Lorenzo, A. (2017). The physiological role of the amyloid precursor protein as an adhesion molecule in the developing nervous system. Journal of Neurochemistry, 143(1), 11–29. 10.1111/jnc.14122

41. Sosa, L. J., Postma, N. L., Estrada-Bernal, A., Hanna, M., Guo, R., Busciglio, J., & Pfenninger, K. H. (2014). Dosage of amyloid precursor protein affects axonal contact guidance in Down syndrome. FASEB Journal, 28(1), 195–205. 10.1096/fj.13-232686

42. Wang, S., Bolós, M., Clark, R., Cullen, C. L., Southam, K. A., Foa, L., Dickson, T. C., & Young, K. M. (2016). Amyloid β precursor protein regulates neuron survival and maturation in the adult mouse brain. Molecular and Cellular Neuroscience, 77, 21–33. 10.1016/j.mcn.2016.09.002

43. Young-Pearse, T. L., Bai, J., Chang, R., Zheng, J. B., Loturco, J. J., & Selkoe, D. J. (2007). A critical function for β-amyloid precursor protein in neuronal migration revealed by in utero RNA interference. Journal of Neuroscience, 27(52). 10.1523/jneurosci.4701-07.2007

44. Zhang, Z., Nadeau, P., Song, W., Donoviel, D., Yuan, M., Bernstein, A., & Yankner, B. A. (2000). Presenilins are required for γ-secretase cleavage of β-APP and transmembrane cleavage of Notch-1. Nature Cell Biology, 2(7). 10.1038/35017108

