## Supplementary material for "APP Dosage and Extracellular Domain Variants Drive Distinct Defects in Neurogenesis modeled in Down Syndrome iPS cells": Figures S1, S2, S3 and S4

### *Supplementary Figures*

#### *Sup. Figure 1*

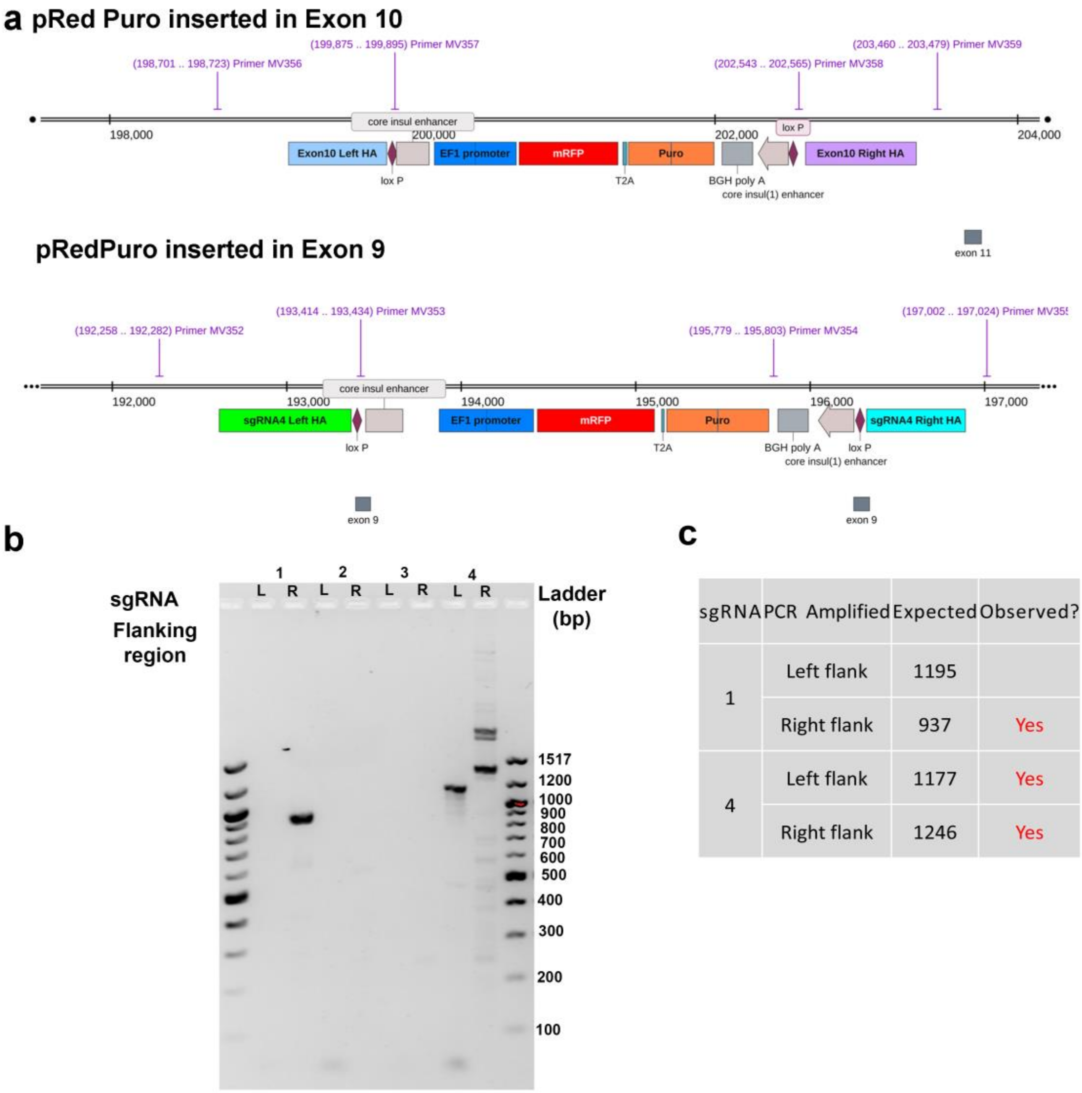


| sgRNA | PCR Amplified | Expected | Observed? |
|---|---|---|---|
| 1 | Left flank | 1195 | |
| | Right flank | 937 | Yes |
| 4 | Left flank | 1177 | Yes |
| | Right flank | 1246 | Yes |

***Figure S1. Validation of pRedPuro integration at the APP locus in modified DS-iPSC lines.***

(a) Schematic representation of the CRISPR/Cas9 strategy showing the targeted integration of the pRedPuro selection cassette into exon 10 (sgRNA 1) or exon 9 (sgRNA 4) of the APP gene. (b) Representative agarose gel electrophoresis of PCR products amplified from genomic DNA using flanking primers that detect the integration of the transgene at the APP locus. Specific bands of expected sizes were observed for sgRNA 1 (APP+/+/Skip10, lane 1) and sgRNA 4 (APP-/-/-, lane 4), confirming successful targeting. (c) Summary table of PCR amplicons showing the expected and observed product sizes, indicating correct transgene integration in both sgRNA target sites.

#### *Sup Figure 2*

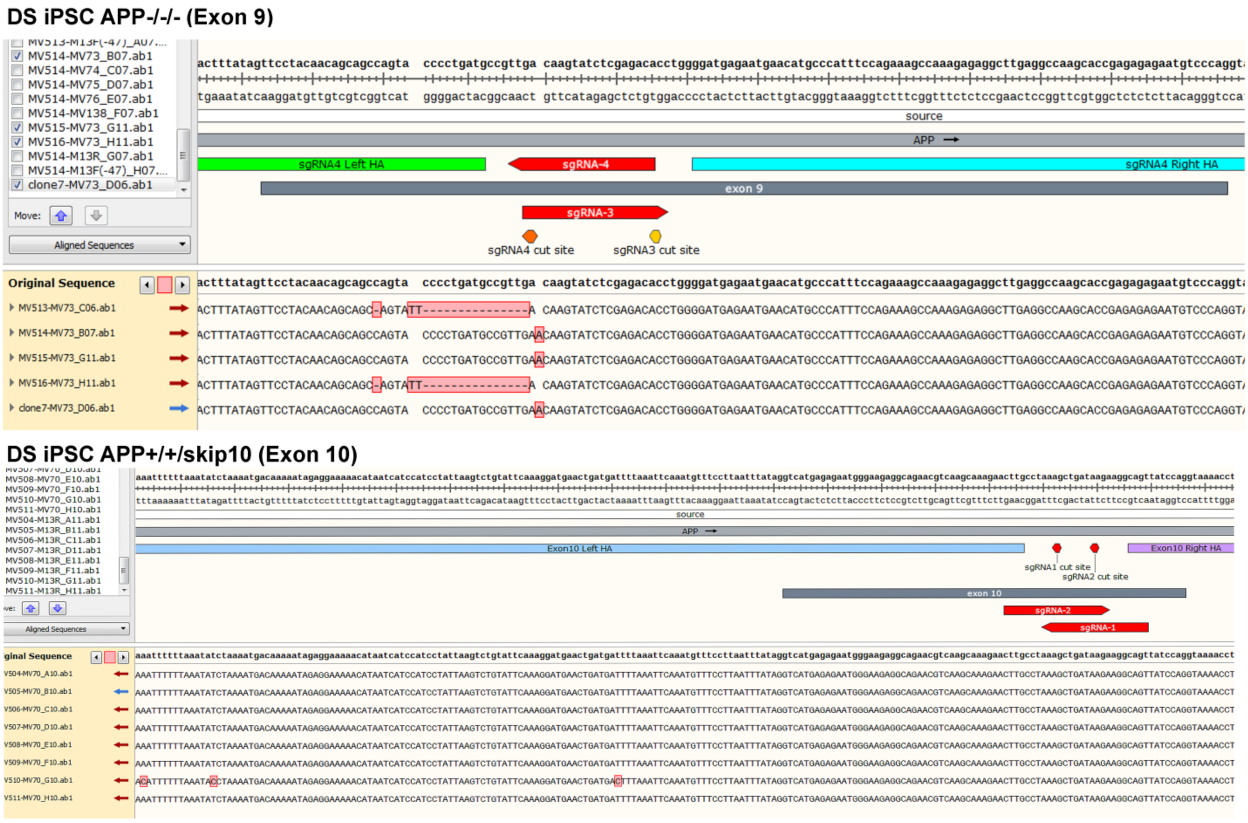


**Figure S2. CRISPR-Cas9–mediated genome editing at the APP locus in DS-iPSC lines.**

Sequencing results following TOPO cloning of PCR products from CRISPR-edited DS-iPSCs. (Top) In APP-/-/- cells (targeted with sgRNA4 at exon 9), sequencing shows insertion-deletion mutations (indels) in the two remaining alleles not carrying the transgene, resulting in biallelic disruption and complete silencing of APP. (Bottom) In APP+/+/Skip10 (sgRNA1 targeting exon 10), no indels were detected in the remaining APP alleles, indicating preservation of two functional copies of APP.

**Sup Figure 3**

**The APP E2 domain**

E2 domain start

LPTTAASTPDAVDKYLETPGDENEHAHFQKAKERLEAKHRERMSQVMREWEEAERQ
AKNLPKADKKAVIQHFQEKVESLEQEAANERQQLVETHMARVEAMLNDRRRLALENY
TALQAVPPRPRHVFNMLKKYVRAEQKDRQHTLKHFEHVRMVDPKKAAQIRSQVMTHL
RVIYERMNQSLSLLYNVPAVAEEIQDEVDEL

**Figure S3.** The sequence of the APP E2 domain (+ 11 AA in front of the domain start), showing locations of sequences encoded by Exon 9 and 10, site of 6nt deletion resulting in change of AVD to D (APP-/-/6nt- cell line), and heparin binding sequences. Exon sequences from UCSC Genome Browser: Human hg38 chr21:25975599-25984058 UCSC Genome Browser v498. E2 Domain and Heparin binding sites derived from UniProt: APP - Amyloid-beta precursor protein - Homo sapiens (Human) | UniProtKB.

Exon 9 (first 11 aa are outside of the reported E2 domain)
AVD - AAs affected by 6 nt deletion. Loss of 6 nt changes AVD to D
Exon 10
Heparin binding sequences

**Sup. Figure 4**

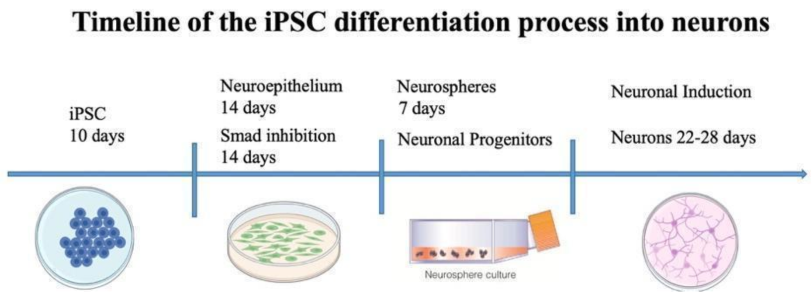


**Figure S4. Stepwise differentiation of DS-iPSCs into neurons.** Schematic representation of the in vitro neuronal differentiation protocol. DS-iPSCs are induced toward neuroepithelial fate via dual SMAD inhibition by day 14, aggregated into neurospheres containing neuronal progenitors by day 21, and subsequently differentiated into neurons between days 22–28. This timeline derives the assays exploring APP dosage effects on neuronal maturation.
